# MARK4 enhances stress granule formation and increases tau accumulation

**DOI:** 10.1101/2025.04.30.651577

**Authors:** Sho Nakajima, Kotone Watanabe, Grigorii Sultanakhmetov, Aoi Fukuchi, Keiya Ito, Sawako Shimizu, Taro Saito, Akiko Asada, Kanae Ando

**Author notes:** Corresponding author Kanae Ando.

## Abstract

Stress granules (SGs) are membrane-less organelles that are formed in response to cellular stress. SG formation protects cells against stress; however, their dysregulation and longer persistence may disrupt proteostasis. Abnormal activity of Microtubule-affinity regulating kinase 4 (MARK4) has been linked to cancer and Alzheimer’s disease. Here, we report that MARK4 is a component of SGs and enhances SG formation. MARK4 is found in SGs in mammalian cultured cells and primary neurons with T-cell intracellular antigen 1 (TIA1). MARK4 expression enhances SG formation induced by hydrogen peroxide. The spacer domain of MARK4 mediates its localization to SGs, and its kinase activity is required for the enhancement of SG formation. MARK4 suppresses TIA1 dimerization induced by hydrogen peroxide. Furthermore, we found that MARK4 and TIA1 synergistically promoted tau accumulation in cultured cells, and knockdown of a fly TIA1 homolog suppressed tau toxicity in a *Drosophila* model. These results indicate that MARK4 regulates TIA1 and SG formation and suggest that MARK4 and TIA1 synergistically contribute to tau toxicity.

## Introduction

SGs are membrane-less organelles composed of untranslated mRNAs, RNA-binding proteins, and various proteins gathered by liquid-liquid phase separation (LLPS) (1). SGs are formed transiently in response to stress and protect cells by interacting with cellular homeostatic processes; however, their persistence has been associated with the pathogenesis of cancer and neurodegenerative diseases (1-3). SG formation in cancer cells enhances tumorigenesis and promotes their adaptation to harsh microenvironments (4). In neurodegenerative diseases that are associated with deposition of protein aggregations, SGs have been suggested to promote conformational changes and abnormal assemblies of disease-causing proteins (5-8).

SGs are heterogenous in their components and properties (9), and SGs containing the RNA binding protein T-cell intracellular antigen 1 (TIA1) have been specifically linked to abnormality of tau protein, whose accumulation causes neuronal loss in a number of neurodegenerative diseases including Alzheimer’s disease (10-14). Hyperphosphorylated tau inclusions colocalize with TIA in cultured cells and postmortem brains of tauopathy patients (10,11). TIA1 promotes SG formations (12) and increases toxic tau oligomers (15), and a reduction in TIA1 protects against neurodegenerative phenotypes in the PS19 tau mice (13,14). TIA1 activity is regulated by oxidation, and reactive oxygen species (ROS) such as H_2_O_2_ oxidize the SG-nucleating protein TIA1 and inhibit SG assembly (3). However, the molecular details underlying the enhancement of tau toxicity via TIA1-containing SGs are not fully elucidated.

Microtubule affinity-regulating kinase (MARK) 4 belongs to the evolutionarily conserved Par-1 family that regulates the cytoskeleton, cell polarity, and energy metabolism (16). Aberrant expression of MARK4 is associated with several types of cancer and with Alzheimer’s disease (17)(18-22). MARK4 phosphorylates tau at Ser262 and Ser356 located in the microtubule-binding repeats (23), and phosphorylation at these sites is critical for tau to gain neurotoxicity (24-29). Mammals express four MARK family members (MARK1–4) with similar functional domains (30,31). Interestingly, although all the members of the MARK family are expressed in the brain and increase tau phosphorylation in the microtubule-binding repeats, only MARK4 has been genetically linked to AD (18-21), and MARK4 enhances tau toxicity more than other members in a *Drosophila* model (32). These reports suggest that upregulation of MARK4 enhances tau toxicity via a MARK4-specific, yet-to-be-identified mechanism.

Here we report that MARK4 is a component of SGs in mammalian cultured cells and primary neurons. Our results suggest that interaction between MARK4 and TIA1 regulate SG formations and MARK4 and tau accumulation. These results suggest a novel function of MARK4 in stress responses and tau abnormality.

## Results

### MARK4 is a component of stress granules

We found that MARK4 shows punctate patterns in the cytoplasm in HeLa cells (Figure 1A). This punctate pattern disappeared when cells were treated with 1,6-Hexanediol, which inhibits LLPS (Figure 1A), suggesting that these puncta are membrane-less organelles. These MARK4-containing puncta were also positive for stress granule markers, TIA1, RasGAP SH3 domain binding protein 1 (G3BP1), or fragile X mental retardation protein (FMRP) (Figure 1B).

**Figure 1.**
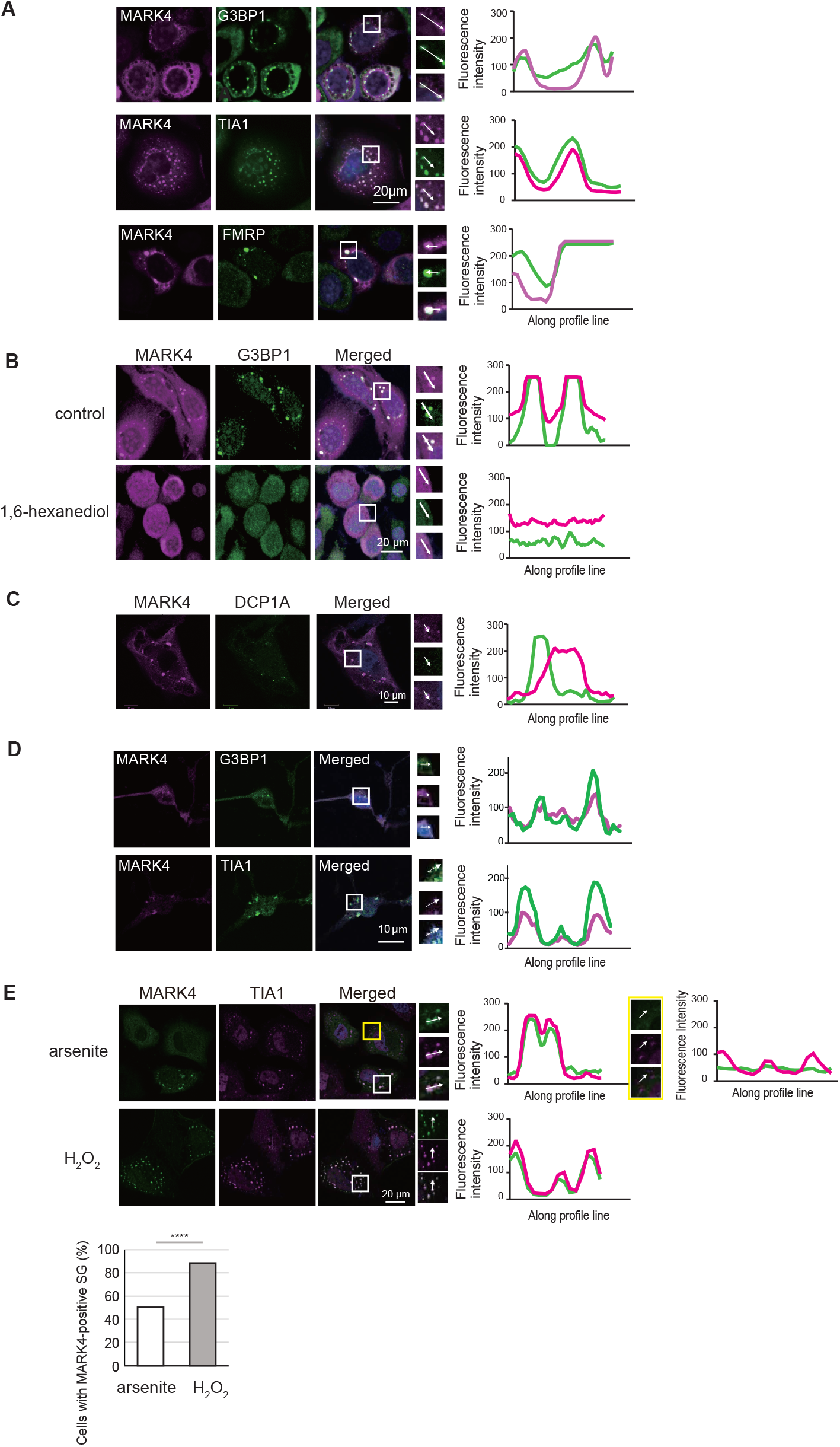
MARK4 localizes to stress granules. (A) HeLa cells were transfected with myc-tagged MARK4 and immunostained with anti-Myc (MARK4, magenta) and antibodies against SG markers (green) such as anti-G3BP1, anti-TIA1 (TIA1) and anti-FMRP. Line plots of fluorescent intensity indicate colocalization of MARK4 and SG markers. (B) MARK4-containing puncta are eliminated by 1,6-hexanediol. Cells were treated with vehicle (top) or 5% 1,6-hexanediol (lower panel) for 3 min and immunostained with anti-Myc and anti-FMRP antibodies. (C) MARK4 does not localize to the p-body. Cells transfected with MARK4 were immunostained with anti-MARK4 and anti-DCP1A antibodies. Representative staining images and line plots. (D) Endogenous MARK4 localizes to SG in primary neurons treated with H_2_O_2_. Mouse primary neurons were treated with H_2_O_2_ and immunostained with anti-MARK4 (Magenta) and anti-G3BP1 (green). (E) MARK4 localizes to stress granules induced by H_2_O_2_. HeLa cells transfected with MARK4 were treated with 0.5 mM sodium arsenite for 1 hr or 1 mM H_2_O_2_ for 1h and immunostained with anti-Myc (MARK4, green) and anti-TIA1 (magenta). TIA-1 positive granules with (white squares) or those without MARK4 (yellow squares) were observed. The percentages of cells SGs containing MARK4 were compared between sodium arsenite-treated cells and H_2_O_2_-treated cells sodium arsenite; n=141, H2O2; n=122; ****, p > 0.001, chi-square test).

Other membrane-free organelles in the cytoplasm formed via LLPS include P-bodies (33). The puncta containing MARK4 were not stained with the antibody against DCP1A, a p-body marker (Figure 1C), suggesting that they are not likely to be p-bodies. These results suggest that MARK4 proteins are localized to stress granules.

We asked whether endogenous MARK4 in neurons localizes to stress granules. Immunostaining of primary neurons that were treated with sodium arsenite to induce stress granule formation revealed that MARK4 co-localizes with stress granule markers (Figure 1D).

### MARK4 preferentially localizes to stress granules induced by oxidative stress

Stress granule formation is induced by various stressors, while the molecular composition and formation mechanisms of SGs differ depending on stress conditions (9). Sodium arsenite and hydrogen peroxide are both known to induce SGs, while hydrogen peroxide induces non-canonical types of SGs (34-37). We analyzed the localization of MARK4 to stress granules formed under different stress conditions. In HeLa cells expressing MARK4, sodium arsenite treatment increased the number of cells with stress granules, and 50% of stress granules contained MARK4. In contrast, in the cells treated with H_2_O_2_, more than 80% of stress granules contain MARK4 (Figure 1E). These results indicate that MARK4 is a component of stress granules induced by H_2_O_2_.

### The spacer domain in MARK4 is required for its localization to stress granules

Mammals express four MARK family members (MARK1–4), and they share the functional domains, such as the kinase domain, UBA domain, spacer domain, and KA1 domain (Figure 2A) (38-40). We asked whether other members of this family localize to stress granules. We found that MARK2 expressed in HeLa cells did not colocalize with G3BP1 (Figure 2A), suggesting that MARK4-specific sequences are required for its localization. Since many proteins localized to stress granules have higher LLPS propensity (41), we compared the LLPS propensity score of MARK2 and MARK4 along their sequence by using *cat*GRANULE (42). We found that MARK4, but not MARK2, has a high LLPS propensity in part of its spacer region (Figure 2B).

**Figure 2.**
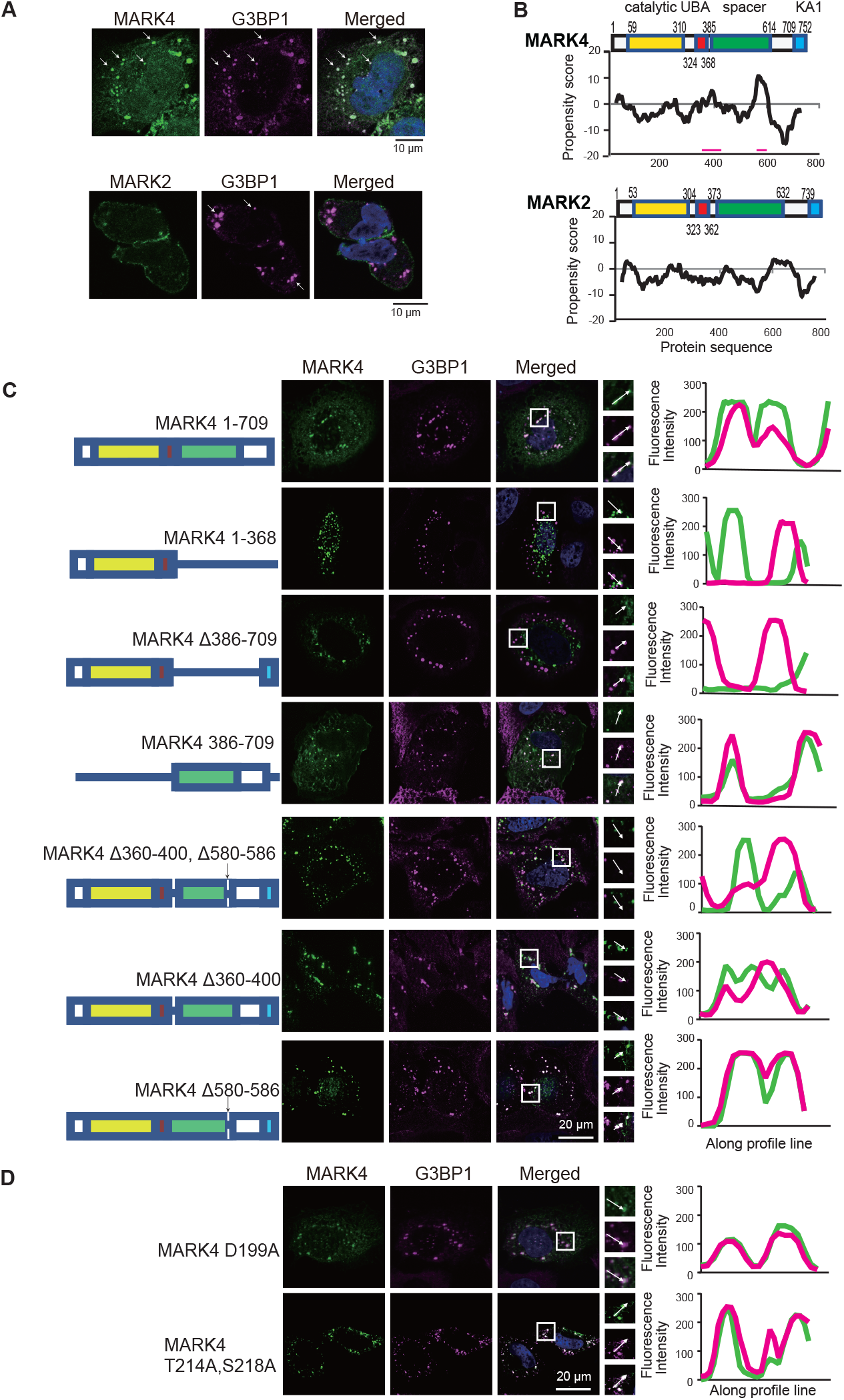
The spacer domain in MARK4 is required for its localization to SGs. MARK2 does not localize to stress granules. Top: Right: HeLa cells transfected with myc-MARK4 or myc-MARK2 were immunostained with anti-Myc and anti-G3BP1 antibodies. Representative images are shown. (B) Schematic diagrams of functional domains and LLPS propensity of MARK2 and MARK4. MARK4 sequence shows higher scores in the spacer domain, especially around 360-400 and 580-586. These sequences are not well conserved between MARK4 and MARK2. (C) Distribution of a series of MARK4 deletion mutants. HeLa cells were transfected with myc-tagged MARK4^1-709^, MARK4^1-368^, MARK4^Δ386-709^, MARK4 ^386-709^, MARK4^Δ360-400,Δ580-586^, MARK4^Δ360-400^, and MARK4^Δ580-586^. Cells were immunostained with anti-Myc (MARK4, magenta) and anti-G3BP1 (green). (D) MARK4 Kinase activity is not required for localization to stress granules. HeLa cells transfected with MARK4 mutants without kinase activity (MARK4^D199A^ and MARK4^T214AS218A^) were immunostained with G3BP1. Representative images and line plots are shown.

To determine the region that mediates the localization of MARK4 to stress granules, we generated a series of deletion mutants of MARK4 and analyzed their colocalization with an SG marker. MARK4^1-709^ colocalized with G3BP1; however, MARK4^1-368^ did not colocalize with G3BP1. Deletion of 386-709 (MARK4^Δ386-709^) abolished its co-localization with G3BP1, while a fragment of MARK4 containing 386-709 (MARK4^386-709^) colocalized with G3BP1 (Figure 2C). These results suggest that MARK4 localizes to stress granules via the spacer domain.

In the spacer domain, MARK4 sequences around 360-400 and 580-586 show peaks in LLPS scores, which are not observed in MARK2 (Figure 2B). We found that the mutant forms of MARK4 with these regions deleted (MARK4^Δ360-400, Δ580-586^) formed foci that did not co-localize with G3BP1 (Figure 2C). We also analyzed the effects of deletion of each region: MARK4^Δ580-586^ co-localized with G3BP1, whereas MARK4^Δ360-400^ formed foci that did not co-localize with G3BP1 (Figure 2C). These results indicate that MARK4^360-400^ is critical for its localization to G3BP1-positive stress granules.

We also examined whether MARK4 kinase activity is required for localization to stress granules. The mutant forms of MARK4 without kinase activity, MARK4^D199A^ (43) and MARK4^T214AS218A^ (44,45), were generated and examined for co-localization with G3BP1. Both mutants co-localized with G3BP1 (Fig. 2D), indicating that MARK4 kinase activity is not required for its localization to stress granules.

### MARK4 promotes stress granule formation

We noticed that MARK4-transfected HeLa cells form SGs more frequently than cells transfected with vectors carrying EGFP, and MARK4-expressing cells contained more SGs compared to cells transfected with EGFP (Figure 3A). We asked whether MARK4 affects the quality of stress granules, such as fluidity, via FRAP analyses of HeLa cells expressing EGFP-TIA1. TIA1 FRAP analyses of HeLa cells expressing EGFP-TIA1 with or without mCherry-MARK4 showed that MARK4 reduces the fluidity of EGFP-TIA1 (Figure 3B).

**Figure 3.**
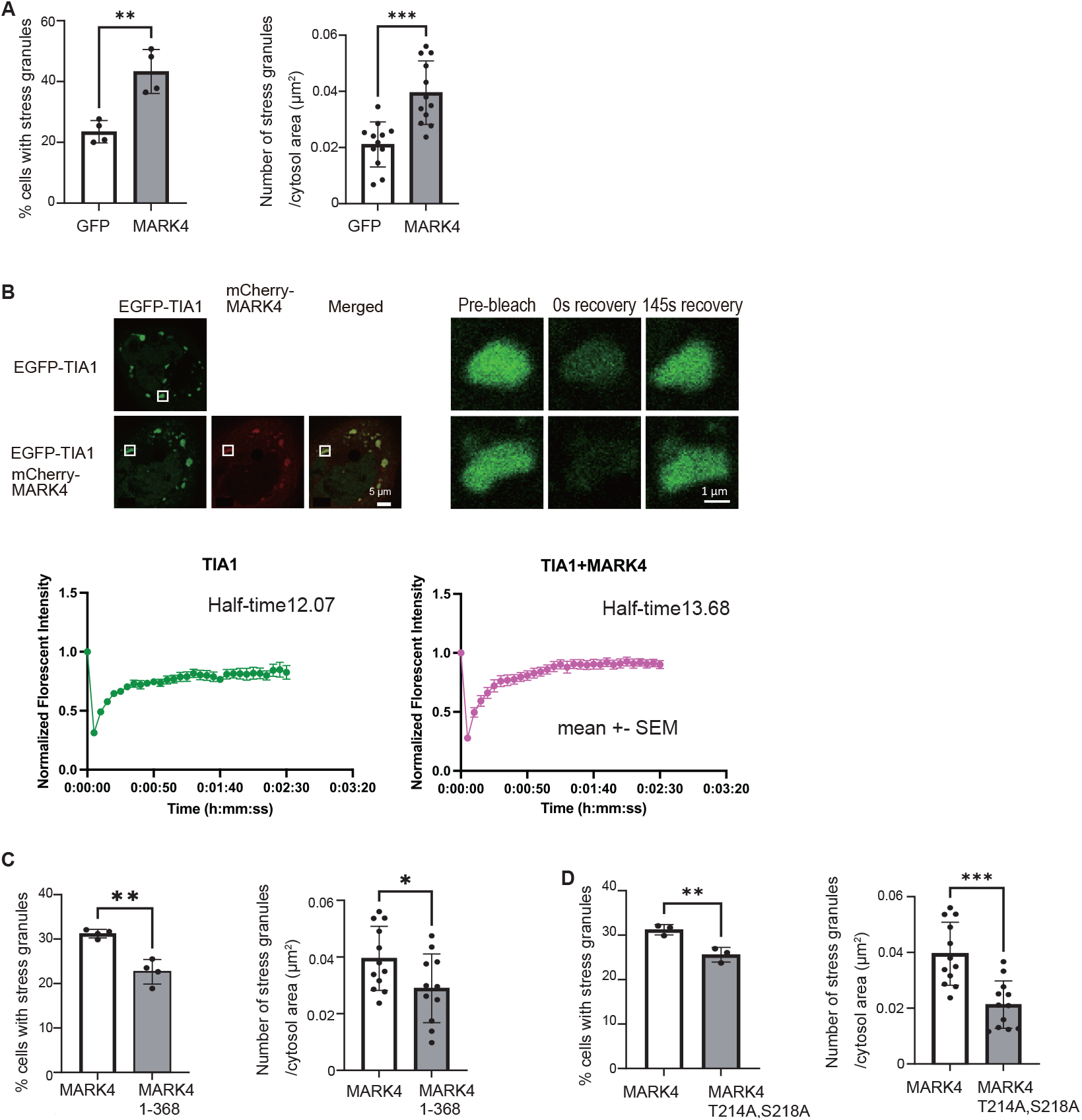
MARK4 promotes stress granule formation. (A) HeLa cells were transfected with EGFP (GFP) or myc-tagged MARK4 (MARK4) and immunostained with anti-G3BP1 to detect SGs. The percentage of SG-containing cells (Mean ± SD, n = 4; **, P < 0.01, Student’s t-test.) and density of SGs (the number of stress granules/cytosol area (mm2)) (Mean ± SD, n = 12; ***, P < 0.005, Student’s t-test.) were higher in MARK4-transfected cells than those in GFP-transfected cells. (B) MARK4 reduces TIA1 fluidity in SG. FRAP was performed in cells transfected with EGFP-TIA1 or EGFP-TIA1 and mCherry-MARK4. Left: Representative images of a SG before the breach, 0 sec after a breach, and 145 sec after a breach. GFP intensity in SGs was normalized to the intensity before the breach and expressed as mean ± standard deviation. ****, P<0.0001 (comparison of fits with <u>extra sum-of-squares F test</u>), best fit values of half-recovery time: 12.07 for TIA1, 13.68 for TIA1+MARK4. (C) MARK4 without SG localization doe not promote SG formation. HeLa cells transfected with MARK4^1-368^ were immunostained with anti-G3BP1 antibodies. The percentage of cells with stress granules (Mean ± SD, n = 3; **, P < 0.01, Student’s t-test.) and the number of SGs per area (Mean ± SD, n = 11-12; **, P < 0. 01, Student’s t-test.). (D) MARK4 without kinase activity does not promote SG formation. HeLa cells transfected with MARK4^T214AS218A^ were immunostained with anti-G3BP1. The percentage of cells with SGs (Mean ± SD, n = 3; **, P < 0.01, Student’s t-test) and the number of SGs per area (Mean ± SD, n = 12; **, P < 0.01, Student’s t-test).

We asked whether MARK4 localization to SGs is required for the enhancement of SG formation. Cells expressing MARK4^Δ1-368^, which does not localize to SGs (Figure 2), had fewer SGs compared to those expressing full-length MARK4 (Figure 3C). Also, cells expressing a kinase-dead form of MARK4 (MARK4^T214AS218A^) showed fewer SGs than those expressing wild-type MARK4 (Figure 3D).

These results suggest that MARK4 localized to SGs enhances SG formation via kinase activity.

### MARK4 reduces TIA1 oxidation

Oxidative stress triggers cellular stress responses to induce SG formation; however, H_2_O_2_ treatment induces oxidation of TIA1 and promotes its dimerization/oligomerization, which reduces its ability to nucleate functional SGs (3). We asked whether MARK4 affects the dimerization of TIA1 induced by H_2_O_2_ treatment. Western blotting revealed that H_2_O_2_ treatment increased the levels of dimerized/oligomerized TIA1 (ox-TIA1) in HeLa cells, as reported in U2OS cells (3). However, cells with MARK4 co-expression showed less ox-TIA1 (Figure 4). This result suggests that MARK4 suppresses TIA1 oxidation and maintains its function to promote SG formation under oxidative stress.

**Figure 4.**
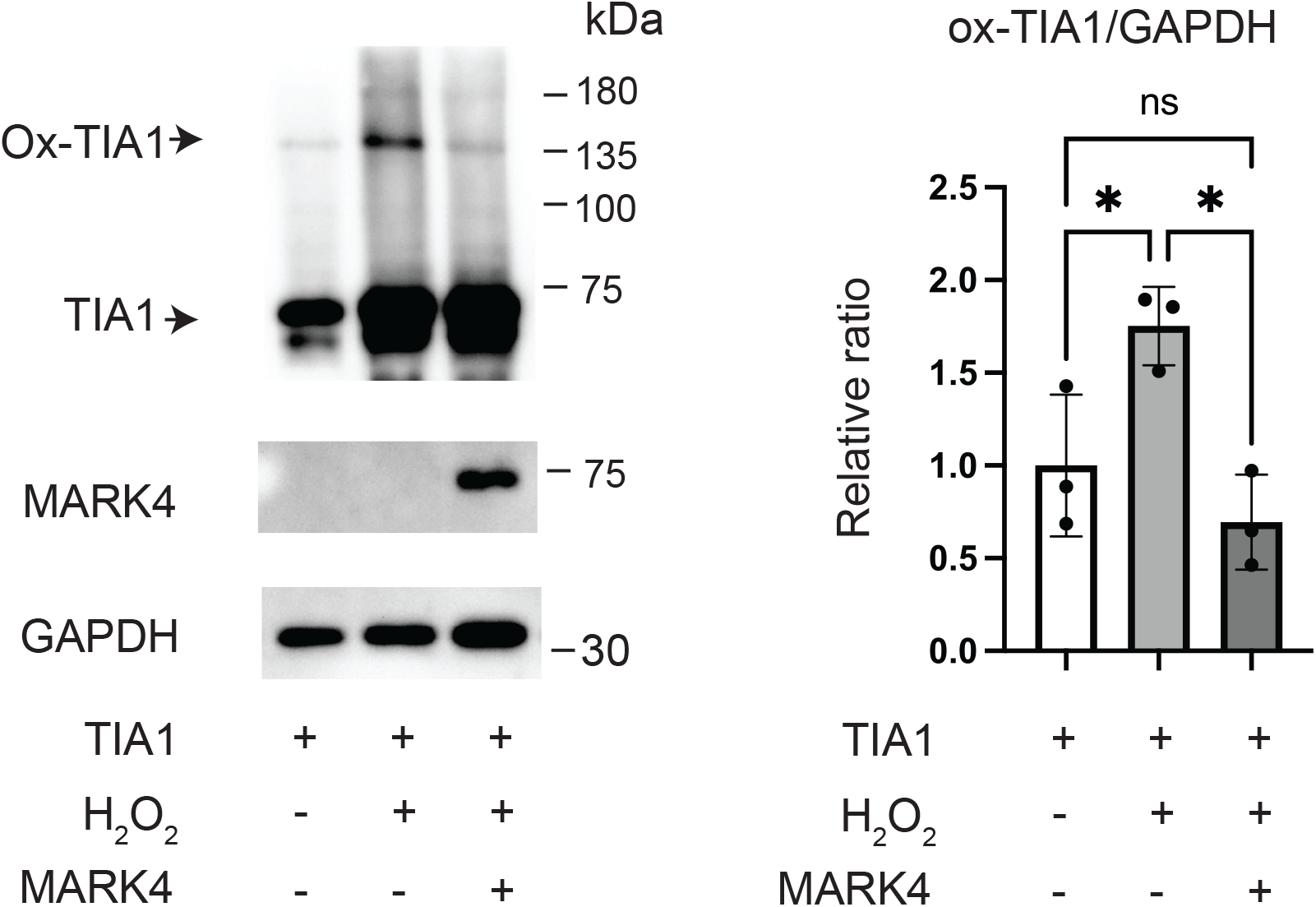
MARK4 reduces TIA1 oxidation. HeLa cells transfected with EGFP-TIA1 or Myc-MARK4 and EGFP-TIA1 were treated with 1 mM H_2_O_2_ for 1h. Cell lysate were subjected to Western blotting with anti-Myc (MARK4) and anti-TIA1. GAPDH was used as a loading control. Representative blots and quantitation (Mean ± SD, n = 3; *, p < 0.05, Ordinary one-way ANOVA tests followed by Tukey’s multiple comparisons test).

### MARK4 and TIA1 synergistically increase tau protein levels

Next, we investigated the effects of MARK4 and SGs on tau proteins. Tau expressed in HeLa cells showed a diffused pattern in the cytoplasm and partially colocalized with MARK4 in TIA1-positive granules (Figure 5A). Western blotting revealed that co-expression of MARK4 and TIA1 increased total tau levels (Figure 5B). Expression of either MARK4 or TIA1 alone did not increase tau levels in HeLa cells (Figure 5B), indicating that the synergistic effects of MARK4 and TIA1 are required for tau accumulation. The levels of Ser262-phosphorylated tau were increased by co-expression of MARK4, and it was further increased by co-expression of MARK4 in TIA1-overexpressing cells (Figure 5B). TIA1 overexpression increased MARK4 levels (Figure 5B), which is likely to cause the increase in pSer262-tau.

**Figure 5.**
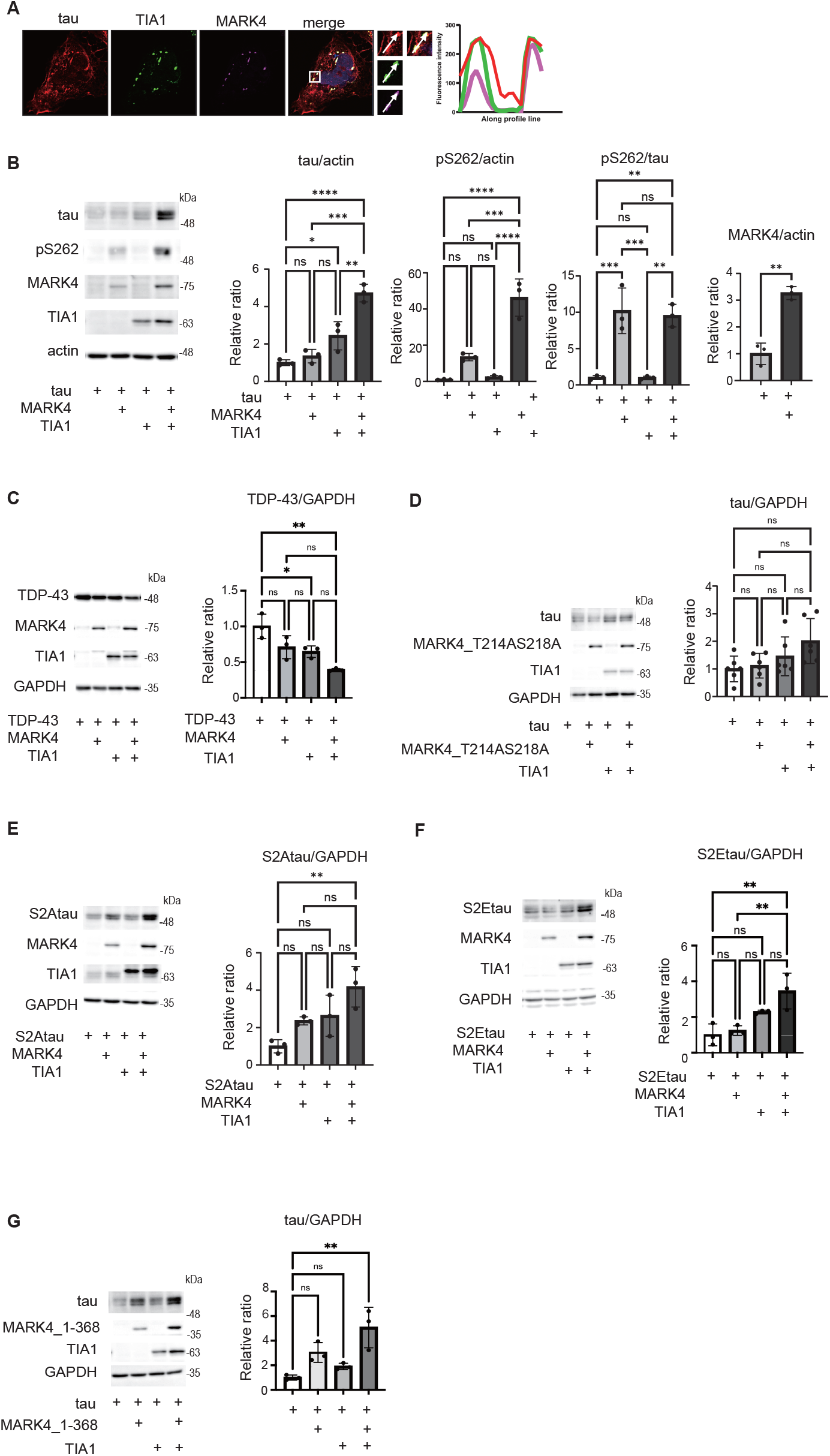
TIA1 and MARK4 synergistically increase tau protein levels. (A) Tau is partially localized to SGs. HeLa cells were transfected with tau and Myc-tagged MARK4 and immunostained with anti-tau (tau, red), anti-Myc (MARK4, magenta), and anti-TIA1 (green). Arrows indicate stress granules containing tau, MARK4, and TIA1. (B) MARK4 and TIA1 overexpression increased levels of total tau and Ser262-phosphorylated tau. Western blotting of HeLa cells transfected with tau, Myc-tagged MARK4, and EGFP-tagged TIA1. Actin was used as a loading control. Representative blot and quantification results are shown (Mean ± SD, n = 3; *, p<0.05, **p < 0.01, ***, p < 0.005, ****, p < 0.0001, Ordinary one-way ANOVA tests followed by Tukey’s multiple comparisons test). (C) MARK4 and TIA1 co-expression does not increase TDP-43. HeLa cells co-transfected with HA-TDP-43 with Myc-MARK4, EGFP-TIA1, or Myc-MARK4 and EGFP-TIA1 were subjected to western blotting. Representative blots and quantitation (Mean ± SD, n = 3; *, p < 0.05, **, p < 0.01, Ordinary one-way ANOVA tests followed by Tukey’s multiple comparisons test). (D) MARK4 without kinase activity does not increase tau levels. HeLa cells were transfected with tau, TIA1, and MARK4^T214AS218A^ and subjected to western blotting. Anti-GAPDH was used as a loading control. Representative blots and quantitative analysis (Mean ± SD, n = 6; ns, p > 0.05, Ordinary one-way ANOVA tests followed by Tukey’s multiple comparisons test). (E) HeLa cells were transfected with tau with alanine substitutions at Ser262 and Ser356 (S2A) and MARK4, TIA1 or MARK4 and TIA1. Cells were subjected to western blotting with anti-tau, anti-Myc (MARK4), and anti-GFP (TIA1). Representative blots and quantification (Mean ± SD, n = 3; **, p < 0.01, Ordinary one-way ANOVA tests followed by Tukey’s multiple comparisons test). (F) HeLa cells were transfected with tau with glutamate substitutions at Ser262 and Ser356 (S2E) and MARK4, TIA1 or MARK4 and TIA1. Cells were subjected to western blotting with anti-tau, anti-Myc (MARK4), and anti-GFP (TIA1). GAPDH was used as a loading control. Representative blots and quantification (Mean ± SD, n = 3; **, p < 0.01, Ordinary one-way ANOVA tests followed by Tukey’s multiple comparisons test). (G) MARK4^1-368^ and TIA1 co-expression increases tau levels. HeLa cells transfected with tau and Myc-tagged MARK4^1-368^, EGFP-tagged TIA1, or Myc-MARK4^1-368^ and EGFP-TIA1, were subjected to Western blotting. using, anti-tau (tau), anti-pS262 (pSer262), anti-Myc (MARK4), anti-GFP (TIA1). Anti-GAPDH was used as a loading control. Representative blots and quantification results (*, p<0.05, **, p < 0.01, Ordinary one-way ANOVA tests followed by Tukey’s multiple comparisons test). (F-G) MARK4 and TIA1 co-expression increased tau with Ser262 substitutions.

We also tested whether the simultaneous expression of MARK4 and TIA1 increases other proteins. TDP-43, which aggregates and accumulates in ALS and FTLD, is known to localize to stress granules (46). We found that MARK4 and TIA1 did not increase TDP-43 levels but rather decreased them (Fig. 5C). These results suggest that MARK4 and TIA1 do not affect protein turnover in general but rather affect tau accumulation via a tau-specific mechanism.

### Tau accumulation caused by TIA1 and MARK4 is independent of tau phosphorylation

Tau phosphorylation at Ser262 and Ser356, the direct targets of MARK4, stabilizes tau proteins and increases their levels in cultured cells, *Drosophila*, and mice (27-29,47). We examined whether MARK4 kinase activity is required for the accumulation of tau protein by MARK4+TIA1. MARK4^T214AS218A^, a mutant lacking kinase activity, did not increase tau phosphorylation at Ser262, as expected (Figure 5D). Co-expression of MARK4^T214AS218A^ and TIA1 did not increase tau protein (Figure 5D), indicating that MARK4 activity is required for tau accumulation in TIA1-overexpressing cells. Unexpectedly, we found that tau phosphorylation at MARK4-target sites was dispensable for its accumulation caused by MARK4 and TIA1: co-expression of MARK4 and TIA1 increased tau proteins with alanine substitutions at Ser262 and Ser356 (S2A) and those with glutamate substitutions at these sites (S2E) (Figure 5E and F), suggesting that co-expression of MARK4 and TIA1 enhances tau accumulation via a mechanism that are independent of phosphorylation of tau at these sites.

We also found that MARK4^1-368^, which does not colocalize to SGs, also increased tau levels when co-expressed with TIA1 (Figure 5G). These results suggest that the accumulation of tau proteins caused by the co-expression of MARK4 and TIA1 is not mediated by direct interaction with tau in SG and suggest that MARK4 and TIA1 synergistically create a cellular condition that favors tau accumulation.

### Knockdown of TIA1 suppresses tau-induced neurodegeneration in *Drosophila*

Finally, we asked how TIA1 is involved in tau-induced neurodegeneration by using a *Drosophila* model. Expression of human tau in *Drosophila* retina induces photoreceptor degeneration. We found that knockdown of Rox8, a functional homolog of TIA1 in *Drosophila* (Figure 6A), suppressed tau-induced retinal degeneration (Figure 6B). Tau protein levels were not significantly affected by knockdown of Rox8 (Figure 6C). Rox8 knockdown also suppressed tau-induced neurodegeneration in the background of MARK4 overexpression (Figure 6D).

**Figure 6.**
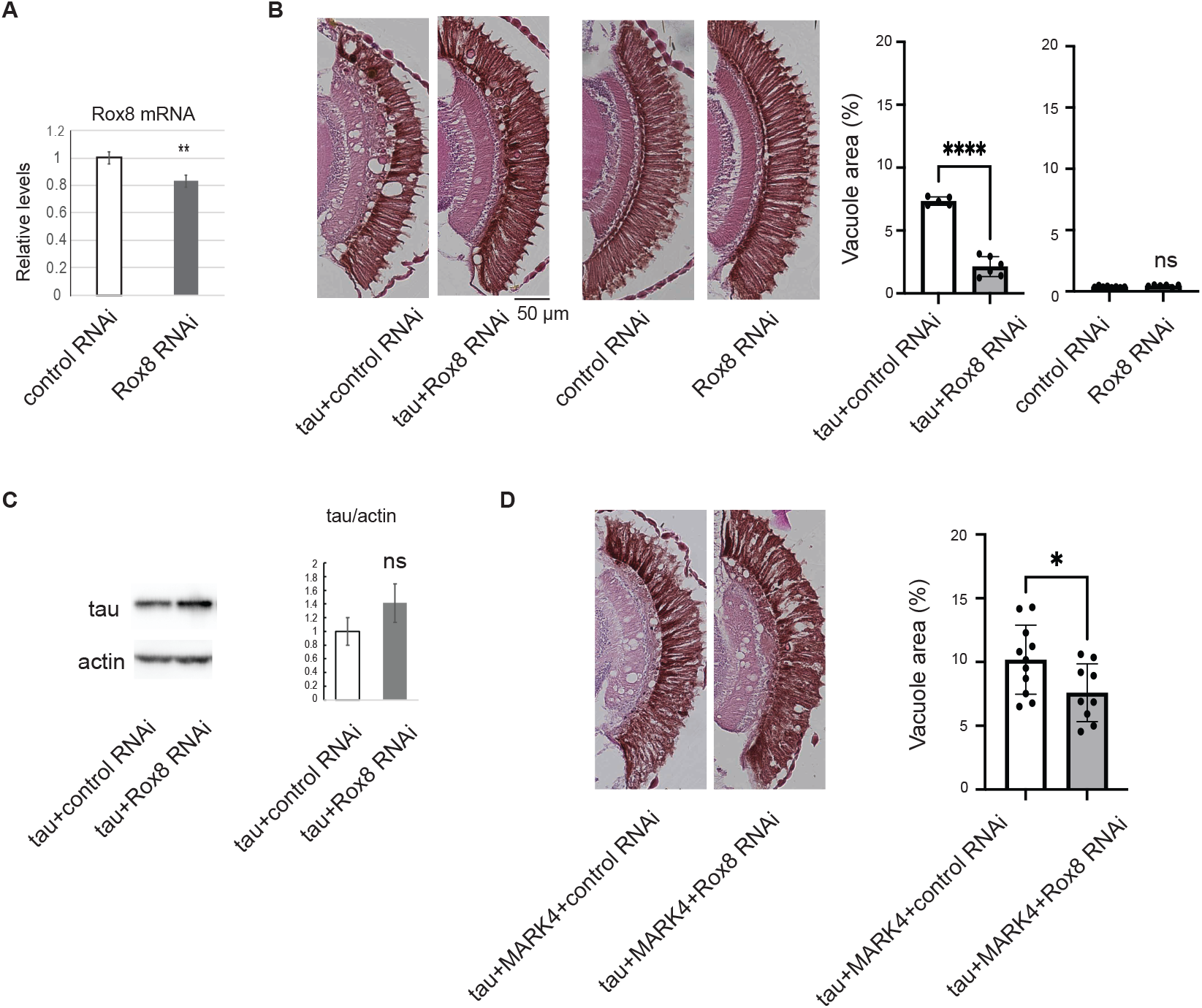
Knockdown of Drosophila homolog of TIA1 suppresses tau-induced neurodegeneration in *Drosophila*. (A) Rox8 RNAi expression reduced Rox8 mRNA. qRT-PCR of head lysate expressing Rox8 RNAi driven by a pan-retinal driver GMR-GAL4. (B) Rox8 expression suppressed tau-induced neurodegeneration. Representative images of paraffin sections of fly retina stained with hematoxylin and eosin. Human wild-type tau was co-expressed with control RNAi (tau+control RNAi) or Rox8 RNAi (tau+Rox8 RNAi) by GMR-GAL4. Without human tau expression, Rox8 RNAi does not cause any morphological change in the retina (compare control RNAi and Rox8 RNAi). Neurodegeneration was identified by the presence of vacuoles (arrows). Scale bar = 20 μm. Vacuole areas in the lamina were quantified and expressed as the ratio (****p<0.0001; Student’ t-test, n = 5). (C) Rox8 knockdown did not affect tau protein expression. Western blotting of head homogenate with anti-tau. Actin was used as a loading control. (n=3, p>0.05, Student’s t-test). (D) Representative images of fly retina co-expressing tau, MARK4, and control RNAi (tau+MARK4+control RNAi) or tau, MARK4, and Rox8 RNAi (tau+MARK4+Rox8 RNAi). Vacuole areas in the lamina were quantified and expressed as the ratio (*p<0.05, n.s., p>0.05, Student’s t-test, n =9). Flies were at ten days post-eclosion.

## Discussion

MARK4 has been reported to play critical roles in the pathogenesis of cancer and neurodegenerative diseases (17-22). MARK4 upregulation in cancer cells enhances metastasis and migration (17), and in the pathogenesis of neurodegenerative diseases, MARK4 phosphorylates microtubule-binding protein tau at the KXGS motif to increase its toxicity (27-29). While MARK family members are highly conserved and all phosphorylate tau at the same sites, MARK4 enhances tau accumulation and toxicity more than other MARKs, suggesting an additional mechanism by which MARK4 enhances tau toxicity (32). In this study, we report that MARK4 is a component of the SG in cultured cells and primary neurons, and a regulator of SG formation in cultured cells under oxidative stress. Since another MARK family member, MARK2, does not distribute to SGs, MARK4 interaction with SGs may explain its specific effects on tau toxicity. Abbarant SG formation has been suggested in the pathogenesis of both cancer and neurodegenerative diseases, and our findings suggest that MARK4 play roles in disease pathogenesis beyond its role as a tau kinase.

Oxidative stress is known as a SG inducer, while its effects are complex (48). Treatment with a high concentration of H_2_O_2_ (1□mM) induces SG formation, while a lower concentration of H_2_O_2_ attenuates arsenite/ER stress-induced SG formation. H_2_O_2_ oxidizes TIA1 and reduces its SG assembly function, thus suppressing SG assembly by impeding the interaction between TIA-1 and its target mRNAs (3). Our results indicate that MARK4 protects TIA1 against H_2_O_2_-induced dimerization, which may contribute to enhanced SG formation under oxidative stress conditions. Supporting this model, MARK4 enhanced SG formation in H_2_O_2_ -treated cells but not in arsenite-treated cells (Figure 1). This result aligns with the involvement of MARK4 in oxidative stress, such as its upregulation in response to oxidative stress and increasing reactive oxygen species in adipocytes (49), cultured cells (50), and a diabetic cardiomyopathy model (51).

We found that knockdown of the *Drosophila* homolog of TIA1, Rox8, suppresses tau toxicity (Figure 6). Tau is an intrinsically disordered protein and is found in multiple assemblies, such as monomers, dimers, oligomers, protofibrils, and fibrils, in diseased brains (52). Once formed, fibrils spread and propagate in a prion-like manner (53). Thus, elucidating the early stage of pathological changes of tau proteins is critical for understanding disease pathogenesis and the development of an effective therapy (54). In *Drosophila*, tau proteins are phosphorylated at disease-related sites and tau neurodegeneration in a phosphorylation-dependent manner without mature fibrils, indicating that this model recapitulates an early stage of tau abnormality (26,55,56). Suppression of tau toxicity in this model by Rox8 knockdown is in line with the reports that SGs affect oligomer formation, the early stage of tau aggregation (57).

Deletion of *Mark4* and reduction of TIA1 both have been reported to reduce tau aggregates and protect against neurodegeneration in the PS19 tau mice (13,14,22), and our findings have revealed the functional link between TIA1 and MARK4 in tau toxicity. MARK4 is a druggable target (58), and its inhibition may be a disease-modifying therapy for cancer and tauopathy.

## Materials & Methods

### Plasmids

Myc-MARK4 and Myc-MARK2 were generated with polymerase chain reaction (PCR)-based methods using the plasmids described previously (59). MARK4 deletion constructs (MARK4^1-709^, MARK4^1-368^, MARK4^Δ386-709^, MARK4^386-709^, MARK4^Δ360-400, Δ580-586^) were generated with Myc-MARK4 by using PCR-based methods. MARK4^D199A^ and MARK4^T214AS218A^ were prepared by using PCR-based site-directed mutagenesis. mCherry-MARK4 was generated with Myc-MARK4 by using Gibson Assembly Cloning Kit (NEB, E5510S). Tau (0N4R) was a kind gift from Dr Mike Hutton (Mayo Clinic Jacksonville). TDP-43 was a kind gift from Dr. Takashi Nonaka (Tokyo Metropolitan Institute of Medical Science) (60). pEGFP-TIA1 is a kind gift from Dr. Mutsuhiro Takekawa (The University of Tokyo) (3).

### Reagent and antibodies

Anti-MYC antibody (4A6, Merck Millipore, 05-724), anti-HA antibody (Proteintech, 51064-2-AP), anti-MARK4 antibody (Proteintech, 20174-1-AP), anti-TIA1 antibody (Proteintech, 12133-2-AP), anti-G3BP1 antibody (Proteintech, 13057-2-AP), anti FMRP antibody (DSHB, 7G1-1), Anti-DCP1A antibody (SIGMA, WH0055802M6), anti-GFP antibody (Merck, 11814460001), anti-tau antibody (T46, Thermo, 13-6400), anti-phosphorylated tau antibody (pS262-tau, Abcam, ab92627), anti-GAPDH antibody(Novus Biologicals, NB100-56875), anti-actin antibody (Proteintech, 66009-1-lg) were purchased. Sodium arsenite (Sigma-Aldrich, 57654633), H_2_O_2_ (Wako, 081-04215), 1,6-hexanediol (Fujifilm Wako, 087-00432) was purchased.

### Cell culture and transfection

HeLa cells were purchased from the RIKEN BioResource Center. Cells were cultured in D-MEM (high glucose) supplemented with 10% deactivated FBS, penicillin, and streptomycin at 37°C in 5% CO2. For immunoblotting and immunocytochemistry, Hela cells were transfected with the indicated plasmids using Lipofectamine 3000 (Invitrogen, Carlsbad, CA) according to the manufacturer’s protocol. Primary neurons were prepared from the mouse brain cortex at embryonic day 15 (E15) and plated on poly-l-lysine-coated dishes in DMEM and Ham’s F-12 (1:1) supplemented with 5% fetal bovine serum, 5% horse serum, 100 U/ml penicillin and 0.1 mg/ml streptomycin at a density of 1.0 × 10^6^ cells/ml. The medium was replaced with Neurobasal medium supplemented with 2% B-27 (Invitrogen), 0.5 mM l-glutamine, 100 U/ml penicillin, and 0.1 mg/ml streptomycin after 4 h of plating.

### Immunofluorescent staining

HeLa cells cultured on glass cover slips were fixed with 4% paraformaldehyde in phosphate-buffered saline (PBS) for 15 min at room temperature. For induction of SG, cells were treated with 0.5 mM Sodium arsenite or 1mM H_2_O_2_ for one hour before fixation. After blocking with PBS containing 5 % skim-milk, cells were incubated with each antibody in PBS containing 5% skim milk and 0.1% Triton X-100 overnight at 4°C followed by incubation with secondary antibodies coupled to Alexa 488 or Alexa 546 (ThermoFIsher, A-11008, A-11001, A-11035, a-11010) (1:500). Nuclei were stained with DAPI. Cells were examined under a fluorescence microscope BZ-700 (Keyence, Osaka, Japan).

### SDS-PAGE and western blotting of cultured cells

HeLa cells were lysed in 20 mM Tris-Cl, pH 8.0, 150 mM NaCl, 1 mM EDTA, 0.1 % SDS, 1 % NP-40, 1 mM dithiothreitol, 10 μg/mL leupeptin, 0.2 mM Pefabloc SC, 0.2mM NaF, and 0.2 mM deoxycholate. After brief centrifugation at 100 g for 5 min, the supernatants were mixed with Laemmli’s SDS-sample buffer containing 0.0625M Tris-HCl (pH6.8), 2% SDS, bromophenol blue, 10% glycerol, and 5% 2-mercaptoethanol. After Laemmli’s SDS–PAGE was carried out, proteins were transferred to PVDF (Millipore) membranes using a semi-dry blotting apparatus or submarine apparatus. Membranes were probed with the primary antibodies, followed by a secondary antibody conjugated with horseradish peroxidase. The reactions were detected using a Millipore Immobilon western chemiluminescent HRP substrate (Millipore).

### FRAP analysis

FRAP was performed with a confocal laser-scanning LSM710 microscope (Carl Zeiss, Jena, Germany) with a 63×/1.4 NA objective lens. Images were acquired every 500 ms. Bleaching was conducted for 100 % laser strength with 50 iterations. GFP intensity of the bleached area was normalized using the intensity of a neighboring cell after background subtraction. Recovery curves were generated and analyzed using Image J and GraphPad Prism10.

### Fly stocks

Flies were maintained at 25°C under light-dark cycles of 12:12 hours, and food vials were changed every 2-3 days. The transgenic fly lines carrying the human 0N4R tau, which has four tubulin-binding domains (R) and no N-terminal insert (N), is a kind gift from Dr. M. B. Feany (Harvard Medical School) (55). GMR-GAL4 and UAS-GFP were obtained from the Bloomington *Drosophila* Stock Center (Indiana University). UAS-MARK4 strain was reported previously (59). UAS-ROX8 RNAi (HMS00472) was obtained from NIG-fly (National Institute of Genetics, Japan). Expression of all proteins was induced by GMR-GAL4 driver in the eyes. All experiments were performed using age-matched female flies.

### Histological analysis

To study the effect of MARKs on tau-induced neurodegenerative phenotype, we analyzed neurodegeneration in the lamina structure ten days after the explosion, as described previously(Iijima et al., 2010). Fly heads were fixated in Bouin’s fixative for 48 h at room temperature, incubated for 24 h in 50 mM Tris/150 mM NaCl, and embedded in paraffin after gradual ethanol dehydration. Horizontal (from top to bottom of the head) 7 μm sections were prepared with standard hematoxylin and eosin staining. Brightfield images of the stained section were captured by the Keyence BZ-X710 microscope. Neurodegeneration in the lamina region was counted as a vacuole area per total lamina size in ImageJ (NIH) using color threshold and particle measure functions. Heads from five to seven flies were analyzed for each genotype.

### Western Blotting of Drosophila heads

Western blotting of Drosophila samples was carried out as described (59). Briefly, 15 heads of flies at 4-5 days after the eclosion per genotype were homogenized in 2xLaemmli Buffer, boiled for 5 min at 95 °C, centrifuged at 15,800 g for 5 min to remove debris, and separated by 10% acrylamide SDS-PAGE. Proteins were transferred to PVDF membranes (Millipore). Membranes were blocked in 5% skim milk in TBS-T and probed with primary antibodies: Anti-Myc Tag Antibody (clone 4A6, Sigma-Aldrich), tau monoclonal antibody (T46, Invitrogen, 13-6400), anti-pS262-tau antibody (Abcam, ab92627), anti-pS356-tau antibody (Abcam, ab92682), anti-actin antibody (Sigma-Aldrich, A2066), anti-pS202/pThr204-tau antibody (AT8, Invitrogen, MN1020), anti-pSer396-tau antibody (Invitrogen, 44-752G). Membranes were incubated with peroxidase-conjugated anti-mouse IgG (DAKO, P0447) and anti-rabbit IgG (DAKO, P0399). Proteins were visualized using Immobilon Western Chemiluminescent HRP Substrate (Millipore) and detected by Fusion FX (Vilber). The integral intensity of bands was quantified by ImageJ (NIH). Western blot experiments were repeated at least three times with independent cohorts.

### qRT-PCR

Quantitative reverse transcription PCR was carried out as previously reported (32). More than 30 flies for each genotype were collected and frozen. Heads were mechanically isolated, and total RNA was extracted using Isogen Reagent (NipponGene, Tokyo, Japan) according to the manufacturer’s protocol with an additional centrifugation step (11,000 g for 5 min) to remove cuticle membranes prior to the addition of chloroform. Total RNA was reverse-transcribed using PrimeScript Master Mix (Takara Bio, Shiga, Japan). qRT-PCR was performed using TOYOBO THUNDERBIRD SYBR qPCR Mix (Osaka, Japan) on a Thermal Cycler Dice Real Time System (Takara Bio). The average threshold cycle value (CT) was calculated from at least three replicates per sample. Expression of genes of interest was standardized relative to rp49. Relative expression values were determined by the ΔΔCT method. Primers were designed using Primer-Blast (NIH) or obtained from Fly Primer Bank (http://www.flyrnai.org/flyprimerbank).The following primers were used for RT-PCR: Drosophila Rox8, forward 5′-AGCCGAAGACAGACATCAGTT -3′, reverse 5′-CTCTCCGAATGGGGCGAAAG -3′, Drosophila rp49, 5′-GCTAAGCTGTCGCACAAATG-3′, reverse 5′-GTTCGATCCGTAACCGATGT-3.

### Statistical analysis

The images of protein domain structure were created in DOG 2.0 (61). Data analysis was performed using Microsoft Excel and GraphPad Prism v9.4. Values bars are given as mean ± standard deviation. Data were analyzed by a one-way ANOVA with Tukey’s or Dunnett’s multiple comparison test. If data did not pass the normality test, data were analyzed by Kruskal–Wallis test with Dunn’s multiple comparisons. p-values <0.05 were considered statistically significant.

## Data availability

The datasets used and analyzed in this study are available from the corresponding author upon request.

## Supporting information

This article contains supporting information.

## Acknowledgement

The authors thank Dr. Mel Feany (Harvard University) for fly stocks, Dr. Mutsuhiro Takekawa (The University of Tokyo) for pEGFP-TIA1 cDNA, and Dr. Takashi Nonaka (Tokyo Metropolitan Institute of Medical Science) for TDP-43 cDNA.

## Ethics declarations

The study was approved by the Research Ethics Committee of Tokyo Metropolitan University.

## Author contributions

Sho Nakajima: Investigation, Formal analysis, Writing – original draft

Kotone Watanabe: Investigation, Validation

Grigorii Sultanakhmetov: Investigation, Formal analysis

Aoi Fukuchi: Investigation

Keiya Ito: Investigation

Sawako Shimizu: Investigation

Taro Saito: Investigation, Resources, Supervision

Akiko Asada: Investigation, Resources, Supervision, Writing – review & editing

Kanae Ando: Conceptualization, Funding acquisition, Investigation, Project administration, Supervision, Writing – original draft, Writing – review & editing.

## Abbreviations

SG: (stress granule)
MARK: (Microtubule-affinity regulating kinase)
TIA1: (T-cell intracellular antigen 1)
LLPS: (liquid-liquid phase separation)
G3BP1: (RasGAP SH3 domain binding protein 1)

